# Enzymatic synthesis of gene-length single-stranded DNA

**DOI:** 10.1101/200550

**Authors:** Rémi Veneziano, Tyson R. Shepherd, Sakul Ratanalert, Leila Bellou, Chaoqun Tao, Mark Bathe

**Author notes:** These authors contributed equally to this work (Rémi Veneziano, Tyson R. Shepherd).

## Abstract

Single-stranded DNA (ssDNA) increases the likelihood of homology directed repair with reduced cellular toxicity, yet ssDNA synthesis strategies are limited by the maximum length attainable, as well control over nucleotide composition. Here, we apply purely enzymatic synthesis to generate ssDNA greater than 15 kb using asymmetric PCR, and illustrate the incorporation of diverse modified nucleotides for therapeutic and imaging applications.

---

Efficient ssDNA synthesis on the 10+ kb-scale is a major need for numerous biotechnology applications including templated homology directed repair for genome editing (1–4), systems-scale gene synthesis and cloning (5–9), and scaffolded DNA origami (10,11). Conventional ssDNA synthesis is performed using either chemical or enzymatic approaches. Chemical synthesis is currently limited to approximately 98% incorporation efficiency for each base addition and therefore limited to the production of ssDNA oligos up to only 200 bases(5). Standard enzymatic synthesis through ligation or polymerization yields double-stranded DNA (dsDNA) that requires additional steps to generate ssDNA. Commercially available ssDNA synthesis is limited to 2 kb from Integrated DNA Technologies, Inc. (IDT, Coralville, IA) or recommended up to 5 kb using a strandase enzyme-based approach from Takara Biosciences, Inc. (Mountain View, CA). Enzymatic or chemical approaches to denaturation of dsDNA to form ssDNA is an alternative approach to ssDNA production, but limited by low yield and high cost.

In contrast, asymmetric polymerase chain reaction (aPCR) offers the direct synthesis of ssDNA from an underlying dsDNA template and has been applied to generate ssDNA ranging from several hundred to several thousand nucleotides in length (12–15). aPCR differs from traditional PCR by having one primer (the forward primer) in molar excess over the second primer (the reverse primer). This approach has previously been applied to short ssDNA synthesis for aptamers and gene detection (12,15), and more recently to kb-scale ssDNA for scaffolded DNA origami (11). However, previous work was limited to 3.3 kb due to low enzyme processivity. Here, we overcome this limitation by using a highly-processive LongAmp Taq polymerase to achieve 15+ kb length ssDNA. Additionally, using a standardized protocol and rules-based primer design, we achieve pure product yields up to 690 ng per 50 μl reaction volume and demonstrate direct incorporation of chemically modified nucleotides for ssDNA applications in therapeutics and imaging that require base or backbone modifications.

High-fidelity polymerases such as Phusion® allow for long dsDNA synthesis in standard PCR, however, Phusion polymerase was unable to synthesize fragment large than 1 kb ssDNA (Fig. 1A **and S1**), likely due to enhanced exonuclease activity (**Fig. S2**). Thus, we evaluated 10 polymerases, representative of different families of enzymes, for their activity in generating ssDNA (Fig. 1A **and S1, Supplementary Table 1**). Based on agarose gel band intensities, *Taq* polymerases gave the highest ssDNA production, possibly due to lower exonuclease activity, with QuantaBio AccuStart HiFi yielded the largest amount of ssDNA with the fewest contaminating dsDNA off-target strands for both 1,000 nt (Fig. 1A, **Line 2 boxed**) and 3,281 nt fragments (**Fig. S1**). To ensure single-strandedness of the product, the reaction was incubated with S1 and *Exo*l nucleases, both specific to ssDNA, showing only digestion of the low molecular weight band (Fig. 1B), while using dsDNA-specific restriction endonucleases (*Eco*RI + *Nae*I) showed digestion of the dsDNA high molecular weight band (Fig. 1B). *Taq* polymerase was capable of synthesizing ssDNA from natural templates such as M13 and from templates synthetic in origin such as from a digital information-encoding sequence (**Fig. S3** and **Supplementary Table 2**). A general protocol was developed for the highest product yield, specific to AccuStart HiFi polymerase, showing 2 mM MgSO_4_ concentration, 1:50 to 1:65 reverse:forward primer ratio, 0.6 ng/μL template concentration, and up to 40 cycles (**Fig. S4 and S5; External Supplementary Excel Table**), with gel extraction being used for subsequent purification (**Fig. S6**) yielding an average synthesis of of 695 ± 35 ng of long ssDNA per 50 μL of aPCR reaction (**Fig. S7**).

**Figure 1.**
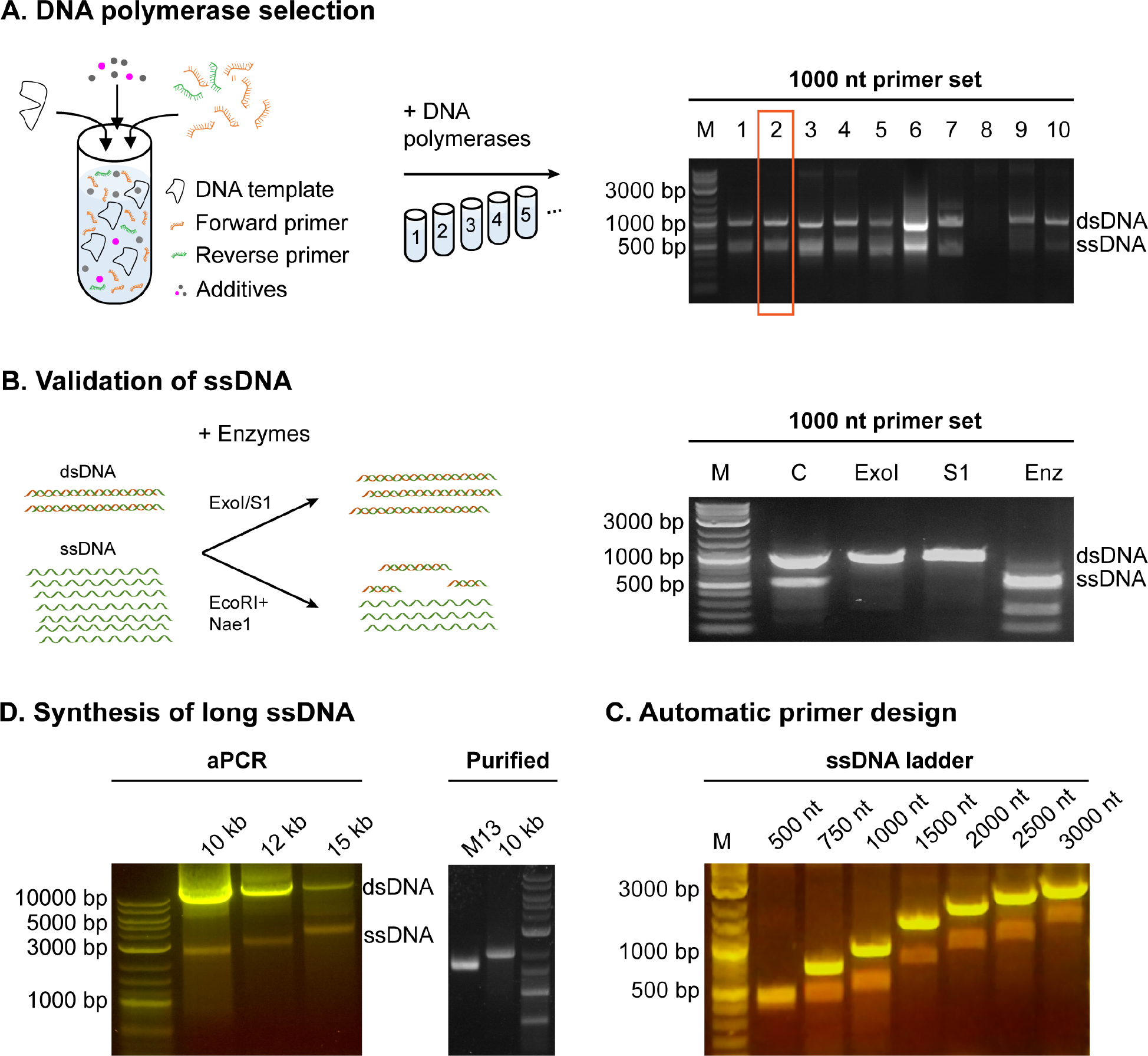
ssDNA production by aPCR. **A.** aPCR reactions were assembled with a 50-molar excess of a forward primer and 10 different polymerases were tested for highest yield of ssDNA (lower bands) as judged by gel electrophoresis (right). QuantaBio AccuStart HiFi, polymerase 2, boxed, produced the highest amount without overlapping dsDNA contaminants. 1. Accustart; 2. Accustart HiFi; 3. Accustart II; 4. AccuPrime; 5. GoTaq; 6. DreamTaq; 7. Phusion; 8. Platinum SuperFi; 9. Q5; 10. Tth polymerase. **B.** *Exo*I and S1 nucleases were used to digest the ssDNA band while *Eco*RI+*Nae*I mix digested the dsDNA band. **C.** An automated primer design algorithm was made to simplify primer selection and validated with product sizes between 500 and 3000 nt. **D.** NEB LongAmp was used to generate ssDNA up to 15,000 nt long.

AccuStart HiFi was capable of synthesizing ssDNA up to 6000 nt, but with reduced yield (**Fig. S8 and Supplementary Table 3**). Initial tests with two other *Taq*-based polymerase sets, NEB LongAmp® *Taq* and Takara LA® *Taq*, produced notable amounts of dsDNA byproduct when tested for amplification of the 1,000 nt and the 3,281 nt fragments (**Fig. S9 and Supplementary Table 4**). However, these byproducts were avoided by increasing the annealing temperature (**Fig. S9**). Given the capacity for these polymerases to synthesize long dsDNA fragments, these enzymes were tested for use in 10+ kb-length ssDNA synthesis. Lambda phage genomic DNA (NEB) was used as a template for long-strand synthesis, with the protocol being only slightly modified, including using less template (0.01 to 0.1 ng/pμL) and increasing the extension time commensurate with the product length. With these modifications, the LongAmp and the LA *Taq* enzymes were capable of producing ssDNA products 10, 12, and 15 kb in length (Fig. 1D and **S10**). While both of the enzymes were capable of synthesis of long fragments, the NEB LongAmp gave the highest yield according to gel band intensity (**Fig. S11**).

To ensure highest yield of user-defined product lengths, primer design rules were generated to reduce off-target sequence amplification, as exponential amplification of undesirable off-target dsDNA will exceed target linear ssDNA production. These include the forward and reverse primers not priming at off-target sequences on the template or product. Additionally, similar to LATE-PCR, the melting temperature of the forward primer should be 1-3°C less than the melting temperature of the reverse primer due to the higher concentration of the former. To reduce mispriming, the forward primer should be more GC-rich in the 5’ half than the 3’ half, and the 3’-nucleotide should terminate in an A or T. Additionally, we found highest ssDNA yield when the forward primer melting temperature is between 54 and 57°C. We codified the aPCR-specific rules into an algorithm for rapid retrieval of application-specific primer sets for user-selected product lengths (named “aPrime”) and experimentally validated the algorithm for products ranging from 500-15,000 nt (Fig 1C **and D and Supplementary Table S5**). Notably, additional template constraints such as limiting high GC-content and avoiding long regions of sequence self-similarly should be avoided.

We extended the capabilities of single-strand synthesis to incorporate modified dNTPs dispersed throughout the entire polymer for therapeutic and imaging applications, similar to what has been shown in dsDNA synthesis (16). We tested this strategy using four different dNTPs replacing in percentage one or all four of the canonical dNTPs. As phosphorothioates are used for nucleic acid polymer stability in the presence of nucleases, we evaluated the efficiency of their incorporation into ssDNA by titrating bulk dNTP phosphorothioate concentration ratios from 0 to 100%. Yield decreased with higher percentage of modified dNTPs to a limit of ~75% before synthesis failed or stalled (Fig. 2A). To test base modification incorporation, we next tested dUTP incorporation into single-stranded DNA synthesis as a replacement for thymidine triphosphate (dTTP). Using pre-generated and purified template generated with 100% dTTP to limit template mutations, we synthesized a gene-length product using complete replacement of dTTP with dUTP (Fig. 2B). For application in molecular coordination, we additionally incorporated biotinylated dNTPs into the synthesized strand (**Fig. S12**). For applications in fluorescence imaging, we evaluated the synthesis of ssDNA with direct incorporation of Cy5-modified dCTP, with up to 10% modified nucleotides (Fig. 2C). Subsequent gel purification and quantification showed up to 2.5% total incorporation on two different templates while using 5% of modified Cy5-dCTP in the dNTP mix (Fig. 2D). The purified fluorescent ssDNA (2000 nt) was used as a scaffold to fold a DNA origami pentagonal bipyramid nanoparticle (Fig. 2E **and Supplementary Table 6**), showing that the chemical modification does not disrupt folding, and can be used for further fluorescent tracking of particles in downstream *in vitro* and *in vivo* biodistribution assays. Additional modification of 10 kb length ssDNA with Cy5-dCTP was also demonstrated, showing long-strand synthesis of fluorescent polymer can be achieved (**Fig S13**). Thus, this work demonstrates the capability of aPCR for production of a variety of useful chemically modified ssDNAs.

**Figure 2.**
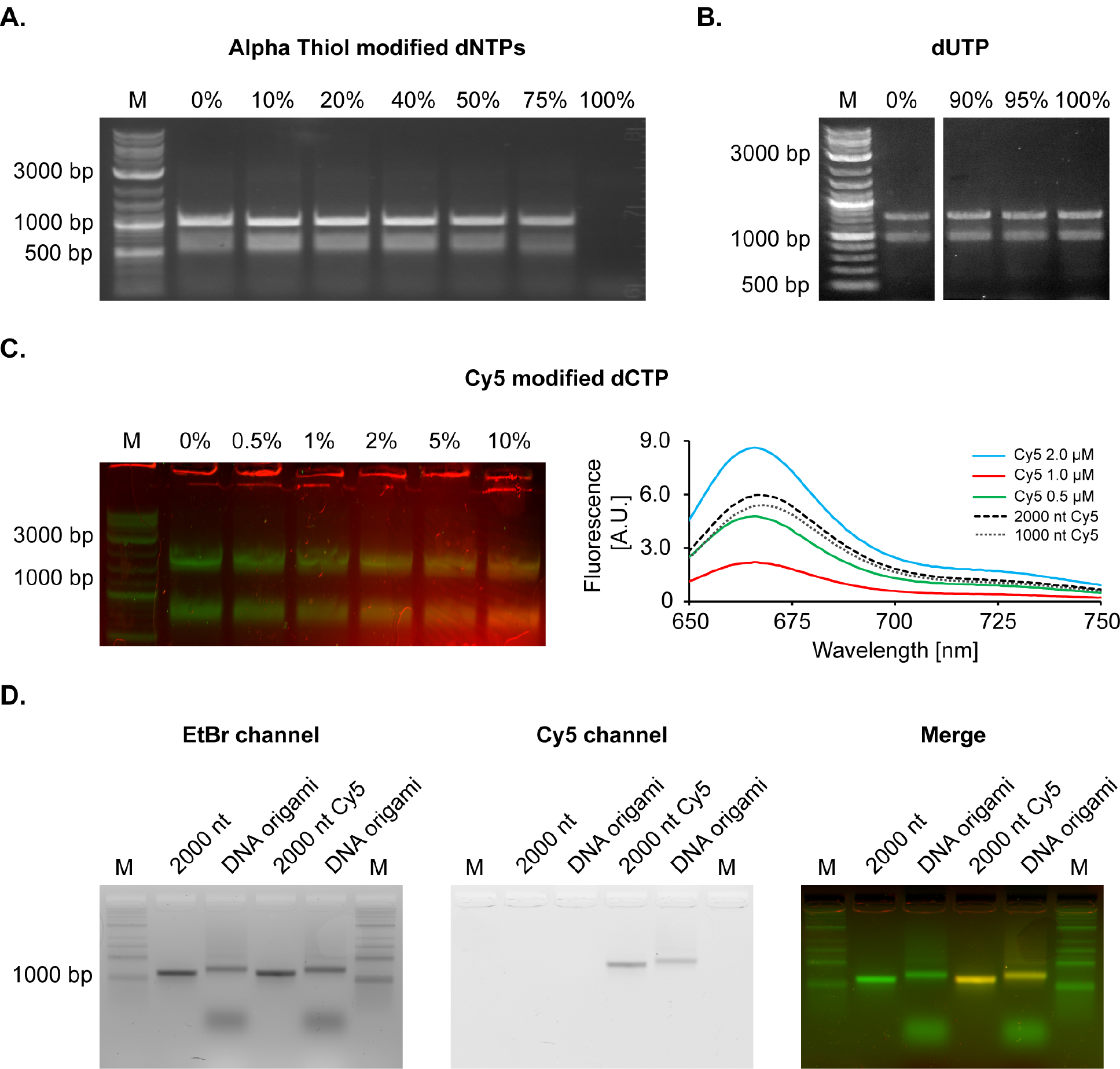
Chemically modified ssDNA produced by aPCR. **A.** Triphosphate-modified nucleotide incorporation by replacing dNTPs with modified nucleotides with incorporation up to 75%. **B.** dUTP replacement for dTTP up to 100% in asymmetric production of the ssDNA. **C.** Cy5 fluorescent modified dCTP incorporated into the synthesized ssDNA, with up to 10% replacement of canonical dCTP (left). Modified ssDNA quantitation showed approximates 2.5% modification of the strand (right). **D.** DNA stain and fluorescent agarose gel mobility shift assay showing folding of a fluorescent scaffolded DNA origami nanoparticle.

In this work, we have extended a simple method for generating ssDNA using aPCR, which allows for synthesis up to 15 kb and additionally allows for one pot chemical modification of ssDNA. The capabilities introduced here will enable future biotechnological applications, including insertion of large chemically- or fluorescently-modified gene constructs through single gene editing experiments, long ssDNA-based memory storage, and scaffold-modified structured nanoparticle synthesis, amongst others.

## METHODS

Methods, including statements of data availability and any associated accession codes and references, are available in the supplementary text. A list of all sequences and primers are contained in the external supplementary Excel table.

## ACKNOWLEDGEMENTS

We acknowledge Dr. William Bricker for aiding in web application development.

## AUTHOR CONTRIBUTIONS

R.V. and M.B. initiated the study; R.V. and T.R.S. designed and analyzed experiments, T.R.S., R.V., and M. B. wrote the manuscript; R.V., T.R.S., and L.B. carried out experiments; T.R.S., S.R., and C.T. programmed aPrime.

